# Influence of Surfactant HLB Values and Agricultural Adjuvants on Pesticide Penetration in Plant Leaves

**DOI:** 10.1101/2025.04.21.649273

**Authors:** Begüm Demirkurt, Maarten Klein, Daniel Bonn

## Abstract

**BACKGROUND:** Effective pesticide action is crucial for optimizing efficacy and minimizing environmental impact, particularly with the increasing reliance on systemic pesticides. Surfactants and adjuvants are commonly used to enhance penetration, but their performance depends on the physicochemical properties of both the pesticide and the surfactants used in the formulations. This study examines how surfactant hydrophilic-lipophilic balance (HLB) values and commercial adjuvants affect pesticide penetration through plant cuticles.

**RESULTS:** We assessed the penetration of two fluorescent pesticide mimics, Rhodamine B (hydrophilic) and Nile Red (lipophilic), into spring onion leaves using confocal laser scanning microscopy. High HLB surfactants significantly enhanced the uptake of Rhodamine B, while low HLB surfactants promoted Nile Red penetration. Surfactants with intermediate HLB values had minimal effect on either compound. Among seven commercial adjuvants tested, only Squall and Prolong significantly improved the penetration of both mimics. Other adjuvants, despite their common use in agriculture, showed limited or no effect on pesticide uptake.

**CONCLUSION:** The HLB value of surfactants strongly influences pesticide penetration, with optimal uptake achieved when the surfactant HLB aligns with the pesticide’s polarity. The penetration mechanisms differ: hydrophilic compounds benefit from increased cuticle hydration with high HLB surfactants, while lipophilic compounds penetrate more effectively with low HLB surfactants that enhance wax fluidity. The limited efficacy of most commercial adjuvants suggests that formulation selection should be based on compound-specific properties rather than generalized claims. These findings emphasize the need for targeted adjuvant selection based on the properties of the active ingredient, providing a practical strategy to optimize pesticide formulations for improved efficacy and reduced environmental impacts.

## 1 Introduction

Pesticides play a crucial role in modern agriculture, helping to control pests and diseases that can significantly reduce crop yields.^1^ Understanding the factors that influence pesticide penetration into plant tissues is essential for optimizing their application and minimizing environmental impact.^2^ Pesticides can be broadly categorized into systemic and contact types based on their mode of action and penetration ability.^3^

Systemic pesticides are absorbed by plants and transported throughout their tissues, including stems, leaves, roots, and flowers.^4^ Unlike contact pesticides that remain on the external surfaces of plants, systemic pesticides utilize the plant’s vascular system for internal distribution. This characteristic allows systemic pesticides to provide long-term protection against pests, even protecting new plant growth.^5^ They are particularly effective against diseases caused by various pathogens, such as insects, fungi, bacteria etc., as these pesticides can spread internally through the plant, offering protection even when new growth occurs or when the pathogen is not directly exposed to the surface.^6^ In contrast, contact pesticides (non-systemic) remain on plant surfaces and are neither absorbed nor dispersed within plant tissues.^7^ Their effectiveness depends on direct contact with target organisms. These pesticides are primarily suitable when immediate control or knockdown of pests is required, but they may not provide long-term protection.^8^ The choice between systemic and contact pesticides depends on various factors, including the target pest, crop type, and environmental considerations.^4^

Among different diseases which need control to protect crop yield, potato late blight infection, a fungal disease caused by *Phytophthora infestans* is agronomically one of the most important.^9^ It is one of the most devastating diseases of potato, and one of the most destructive plant diseases across all major crops. Traditionally, multi-site contact fungicides have been used to control the disease. Mancozeb, long favored for its affordability and broad-spectrum activity, has been banned in the European Union due to concerns over its environmental impact and potential health risks. As a result, there is a shift towards singlesite systemic fungicides, which offer more targeted activity and improved performance under modern application constraints. Unlike contact fungicides, the systematic ones penetrate into the plant tissues, offering extended protection even in untreated areas, and are less prone to wash-off by rain.^9^

Moreover, regulatory changes in pesticide application have introduced new challenges. Over the last decade, the use of anti-drift nozzles has become mandatory in many agricultural settings.^10^ These nozzles, designed to reduce pesticide drift, produce larger droplets, which results in reduced spray coverage. For example, while traditional flat-fan nozzles generate approximately 300 droplets per square centimeter, the 90% anti-drift nozzles only produce about 30 droplets per square centimeter.^10^ This reduction significantly hampers the efficacy of contact fungicides, which rely on thorough surface coverage to be effective. In contrast, systemic fungicides are less dependent on droplet density, as their ability to penetrate plant tissues and spread internally means they can maintain efficacy even under reduced coverage conditions.

While the systemic fungicides deliver effective disease control, they come at a higher cost. The higher price of single-site systemic fungicides contributes to increased crop protection expenses, underscoring the need for strategies that can enhance their penetration performance. In this context, adjuvants, especially surfactants, have emerged as vital tools. By improving the penetration of systemic fungicides into plant leaves, adjuvants can significantly boost the efficacy of systemic products, helping to reduce the required dose and improve overall application efficiency. Some commercial agricultural adjuvants containing surfactants, like Squall, have been shown to improve the performance of crop protection products by helping active ingredients reach their targets more effectively and enhancing rainfastness.^11^

Penetration of pesticides into plant leaves is influenced by numerous factors, including the physicochemical properties of the pesticide, the characteristics of the plant cuticle, and environmental conditions.^2^ The plant cuticle, a waxy layer on the leaf surface, serves as a significant barrier to pesticide penetration.^12,13^ Several techniques have been developed to assess pesticide penetration, including Surface-Enhanced Raman Scattering (SERS) mapping, which allows for in situ and real-time tracking of pesticides using gold nanoparticles as probes.^14^ Studies using SERS have shown that systemic pesticides penetrate more rapidly and deeply into live leaves compared to harvested leaves, with the systemic pesticide thiabendazole reaching depths of 225 *µm* in live leaves after 48 hours of exposure.^14^

Adjuvants, particularly surfactants, play a critical role in enhancing pesticide efficacy by improving their penetration into plant tissues.^15,16^ Surfactants are classified based on their hydrophilic-lipophilic balance (HLB) value, which influences their interaction with different types of pesticides and plant surfaces.^17^ Previous research has demonstrated that hydrophilic surfactants with high HLB values are most effective at enhancing the penetration of herbicides with high water solubility, whereas lipophilic surfactants with low HLB values are better suited for herbicides with low water solubility.^18,19^ The mechanisms by which surfactants enhance pesticide penetration differ based on their HLB values. High HLB surfactants are absorbed into the cuticle and enhance the water-holding capacity (hydration state) of the cuticle, which increases the permeance of hydrophilic pesticides.^18,20^ In contrast, low HLB surfactants increase the fluidity of waxes in the cuticle, as measured by a small reduction in melting point, which enhances the permeance of lipophilic pesticides.^18^

While previous studies have examined the effects of surfactants on pesticide penetration,^21-23^ a systematic investigation using a wide range of HLB values with both hydrophilic and lipophilic pesticide mimics, coupled with an assessment of commercial adjuvants, has not been extensively documented in the literature. This study aims to fill this gap by investigating how surfactants with varying HLB values and commercial adjuvants affect the penetration of hydrophilic and lipophilic pesticide mimics into spring onion leaves.

## 2 Materials and Methods

### 2.1 Pesticide Mimics

Two fluorescent pesticide mimics were used in this study: Rhodamine B (hydrophilic) dissolved in water, and Nile Red (lipophilic) dissolved in a 50% (v/v) water/isopropanol mixture. These fluorescent dyes served as models for hydrophilic and lipophilic pesticides, respectively, allowing visualization of their penetration using confocal microscopy.^21-23^

### 2.2 Surfactants and Adjuvants

Five surfactants with systematically varying HLB values ranging from approximately 2 to 22 were selected to evaluate their effects on pesticide mimic penetration.^17^ The studied surfactants are Tween 20 (T20), Tween 80 (T80), BrijO10 (BO10), Span 20 (S20) and Span 80 (S80). In addition, the following commercial adjuvants commonly used in Dutch agriculture were tested: Prolong (P), Elasto (E), Wetcit Neo (W), Designer (D), Coda Cide (C), Well Power (WP) and Squall (Sq).

### 2.3 Plant Material

Spring onions were chosen for this study due to their waxy, difficult-to-wet leaves, which provide a challenging model for assessing pesticide penetration.^13^ The use of plants with such characteristics allows for a more rigorous evaluation of surfactant and adjuvant performance.

### 2.4 Penetration Assessment Using Confocal Microscopy

A drop of each surfactant or adjuvant solution containing the fluorescent pesticide mimic was applied to the surface of spring onion leaves. After a 24-hour incubation period, excess solution was meticulously cleaned by using cotton buds wetted with 50% (v/v) water/acetone solution, and confocal laser scanning microscopy (CLSM) was employed to assess penetration depth from the leaf surface inward (from adaxial surface to abaxial).^24^ This approach allows for direct visualization of pesticide penetration, similar to methods used in previous studies investigating pesticide behavior in plant tissues.^23^ While SERS mapping using gold nanoparticles has been used for in situ and real-time tracking of pesticides,^14^ our study employed CLSM due to its widespread availability and suitability for fluorescent compounds.^21-23^

Fluorescence imaging was performed using a Leica DMi8 Inverted Confocal Scanning Laser Microscope equipped with a photomultiplier tube (PMT) detector. A 40× air objective (N PLAN L, NA 0.55, Leica) was used to acquire the images. The excitation wavelength was 552 nm for both dyes, and fluorescence emission was collected within the 575–700 nm spectral window. All leaf samples were scanned in the z-direction over 300-400 µm (depending on the sample) with 1.3 µm step sizes for assessing the penetration/diffusion of the pesticide mimics. Cross-sectional images were aligned side by side using the glass surface as the reference plane. Imaging parameters were optimized based on the sample with the lowest fluorescence response. To ensure comparability across different samples, the same imaging settings were used for all samples stained with the same dye, even if this led to saturation in some images.

All image processing and data analysis were performed using ImageJ and Python. Images were despeckled in ImageJ, and the FIRE LUT was applied to enhance contrast for better visualization of intensity variations. Mean diffusion length values were calcuated from thresholded cross-section images. Global Otsu thresholding was applied, and the resulting binarized images were analyzed using a Python script. For each x-coordinate in the cross-section images, the script identified the last z-coordinate with a value of 1, then averaged these values to determine the mean diffusion length for each sample.

### 2.5 Alternative Methods for Assessing Pesticide Penetration

Several other methods have been developed to assess pesticide penetration into plant tissues. The Franz diffusion cell system, as used in a study on apple peel barrier effects, provides quantitative data on pesticide permeation.^25^ This method involves placing plant tissue between donor and receptor compartments and measuring the amount of pesticide that diffuses through the tissue over time.

Liquid chromatography tandem mass spectrometry (LC-MS/MS) can also be used to quantify pesticide penetration through a “spot and wash” approach, where pesticides are applied to leaf surfaces, and the amount remaining after washing is measured.^26^

## 3 Result

### 3.1 Effect of Surfactant HLB Values

The CLSM images revealed distinct patterns of penetration for the hydrophilic and lipophilic pesticide mimics in the presence of surfactants with varying HLB values. High HLB surfactants (values approximately 15-22) significantly enhanced the penetration of the hydrophilic mimic Rhodamine B, while low HLB surfactants (values approximately 2-8) improved the penetration of the lipophilic mimic Nile Red.^18,19,22^

**Figure 1:**
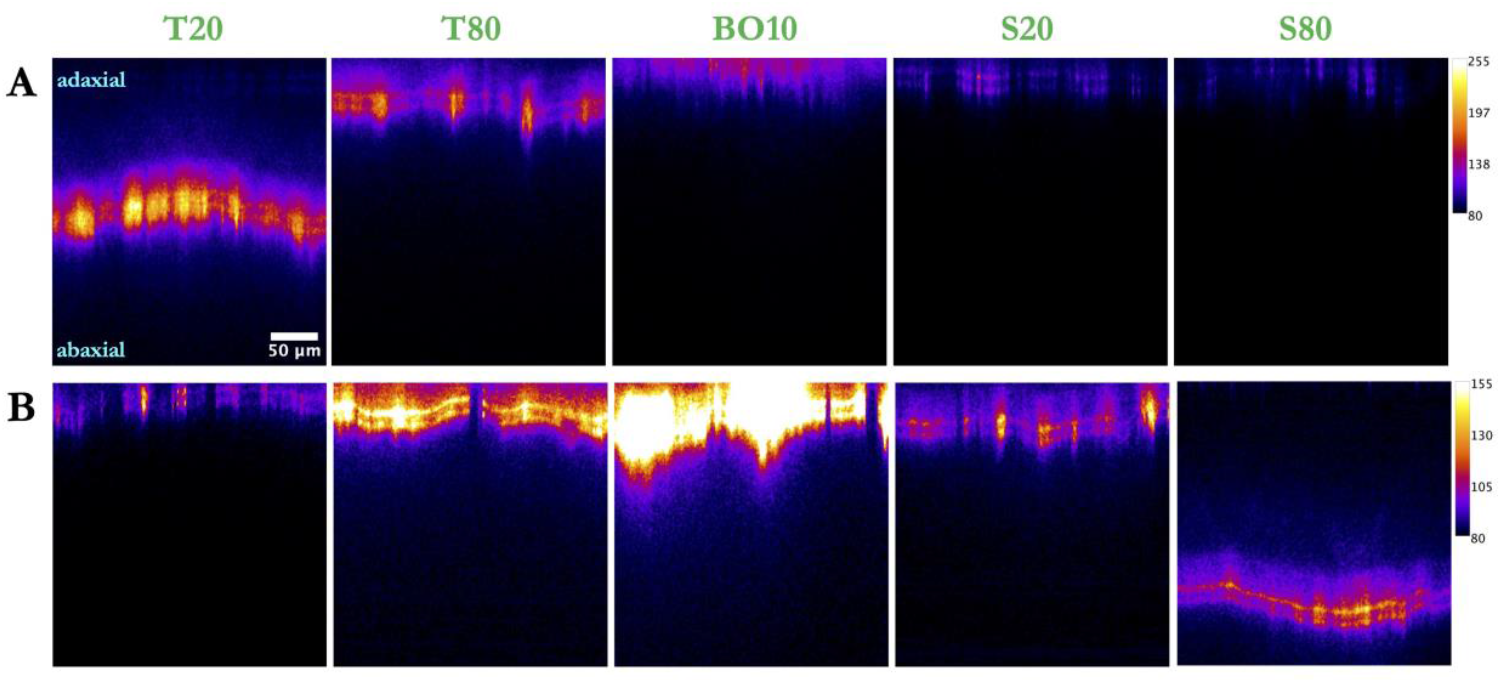
Penetration of Rhodamine B (A) and Nile Red (B) through spring onion leaves in the presence of different surfactants, 24 hours after droplet application. All images show fluorescence intensity cross-sections of the treated areas.

This observation aligns with previous research indicating that hydrophilic surfactants with high HLB values are most effective at enhancing the penetration of water-soluble compounds, whereas lipophilic surfactants with low HLB values are better suited for compounds with low water solubility.^15,18^ The enhancement of pesticide penetration by surfactants with extreme HLB values (either very high or very low) suggests that different mechanisms are involved in facilitating the penetration of hydrophilic and lipophilic compounds through the plant cuticle.^18,20^

**Figure 2:**
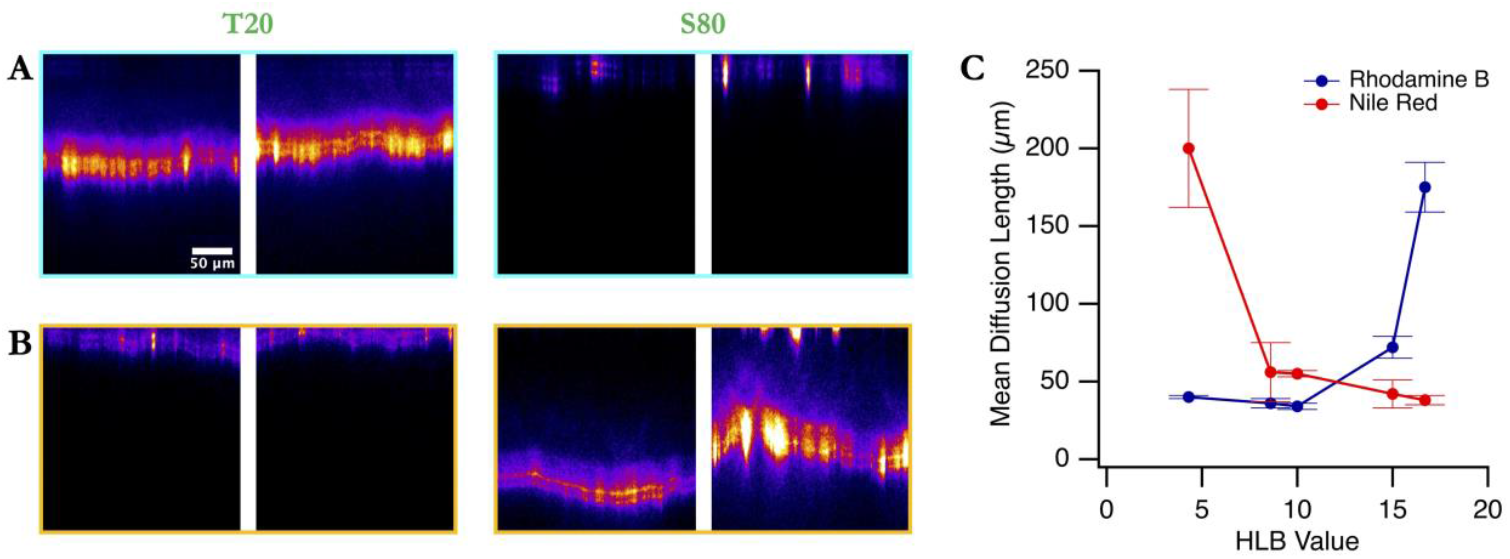
Fluorescence intensity cross-section images showing the penetration of Rhodamine B (A) and Nile Red (B) through spring onion leaves in the presence of Tween 20 and Span 80 surfactants, representing the two extremes of the HLB values studied, 24 hours after droplet application. Images are taken across different areas of the treated leaves. (C) Mean diffusion lengths plotted against the HLB values of the surfactants added to the fluorescent mimic solutions. Standard deviations represent measurements from different areas within the same samples and between different samples.

For Rhodamine B, the penetration depth increased with increasing HLB value of the surfactant, with the highest penetration observed with surfactants having HLB values above 15. In contrast, Nile Red penetration was most enhanced by surfactants with HLB values below 8. Surfactants with intermediate HLB values (9-14) showed limited enhancement of penetration for both mimics.

### 3.2 Effect of Commercial Adjuvants

Among the commercial adjuvants tested, only Squall and Prolong demonstrated significant enhancement of pesticide mimic penetration. Other adjuvants did not significantly affect the penetration of either Rhodamine B or Nile Red into spring onion leaves

The effectiveness of Squall aligns with its documented ability to improve the performance of crop protection products by helping active ingredients reach their targets more effectively. Squall has been shown to enhance rainfastness, with scientific field tests confirming that 50% more active ingredients remain on the crop after a standard rain shower when Squall is added to the tank mix.^27^

**Figure 3:**
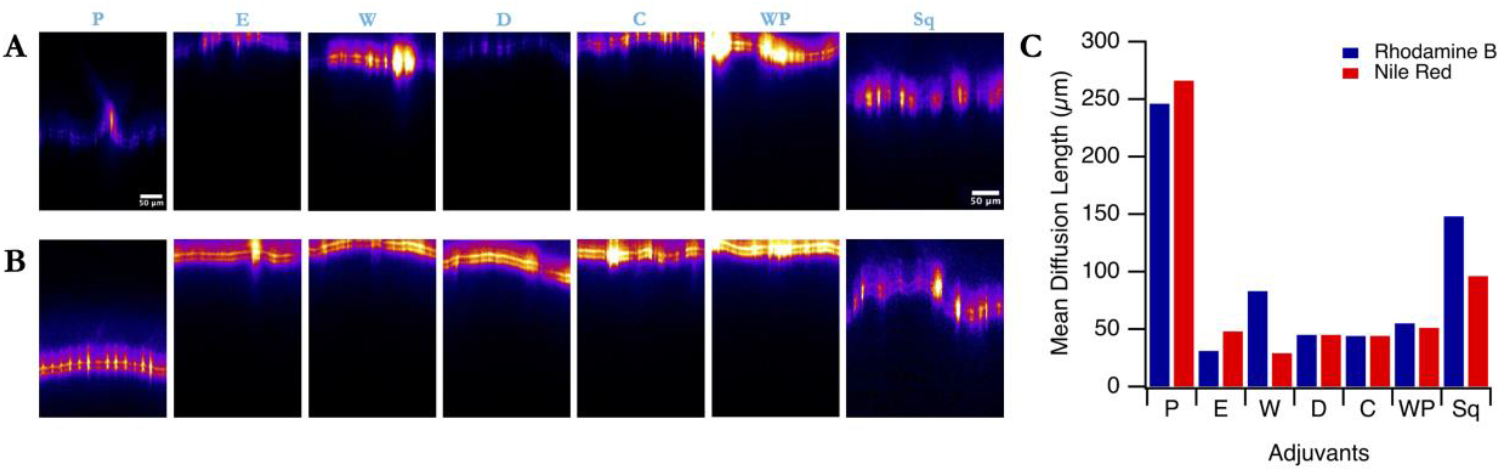
Fluorescence intensity cross-section images showing the penetration of Rhodamine B (A) and Nile Red (B) through spring onion leaves in the presence of commercial adjuvants, 24 hours after droplet application. (C) Mean diffusion lengths plotted against the different adjuvants added to the fluorescent mimic solutions.

The limited effectiveness of most commercial adjuvants in enhancing pesticide penetration suggests that while they may improve spray coverage and retention, they may not necessarily facilitate the movement of pesticides through the plant cuticle. This finding highlights the importance of selecting adjuvants based on specific application requirements, rather than assuming that all adjuvants will enhance penetration.^20^

## 4 Discussion

The results of this study provide important insights into the factors that influence pesticide penetration into plant leaves and the role of surfactants and adjuvants in this process. The observation that high HLB surfactants enhance the penetration of hydrophilic compounds while low HLB surfactants improve the penetration of lipophilic compounds is consistent with the mechanisms proposed in previous research.^18^

High HLB surfactants are believed to enhance the water-holding capacity (hydration state) of the cuticle, which increases the permeance of hydrophilic compounds into the cuticle and subsequently increases their diffusion rate at a constant concentration gradient.In contrast, low HLB surfactants increase the fluidity of waxes in the cuticle, which enhances the permeance of lipophilic compounds.

The effectiveness of Squall and Prolong in enhancing pesticide penetration suggests that these adjuvants either have formulations with extreme HLB values or may incorporate other mechanisms that facilitate pesticide movement through the plant cuticle. The limited effectiveness of most other commercial adjuvants highlights the need for more targeted approaches to adjuvant selection based on the physicochemical properties of the pesticide and the desired mode of action.

This study also demonstrates the utility of confocal microscopy for assessing pesticide penetration.^21-23^ While other methods, such as SERS mapping^14^ and Franz diffusion cells^25^, have been used in previous studies, CLSM provides a direct and visual approach to tracking pesticide movement through plant tissues. This method is particularly useful for fluorescent compounds, such as the pesticide mimics used in this study.^21^

The choice of spring onions with their waxy, difficult-to-wet leaves as a model system provides valuable insights into how surfactants and adjuvants interact with challenging leaf surfaces.^13^ Waxy cuticles present a significant barrier to pesticide penetration, and understanding how to overcome this barrier is crucial for optimizing pesticide application strategies

In the broader context of pesticide use in agriculture, the findings of this study have important implications for the ongoing shift from contact to systemic pesticides. Systemic pesticides have been favored for their ability to provide long-term protection by being absorbed and distributed throughout the plant. However, concerns about their environmental impact, particularly on non-target organisms, have led to increased scrutiny. Enhancing the penetration of pesticides through the strategic use of surfactants and adjuvants could potentially allow for lower application rates, thereby reducing environmental impact while maintaining efficacy.

## 5 Conclusion

This study demonstrates that the HLB value of surfactants plays a critical role in determining their effectiveness in enhancing pesticide penetration into plant leaves. High HLB surfactants significantly improve the penetration of hydrophilic compounds, while low HLB surfactants enhance the penetration of lipophilic compounds. Among commercial adjuvants, only Squall and Prolong showed significant enhancement of pesticide penetration, highlighting the need for careful selection of adjuvants based on specific application requirements.

The findings of this study contribute to our understanding of how surfactants and adjuvants influence pesticide behavior in plant systems and provide valuable insights for optimizing pesticide formulations and application strategies. Future research should focus on further elucidating the mechanisms by which surfactants and adjuvants enhance pesticide penetration and on developing more effective formulations that balance efficacy with environmental safety.

